# Using endocytosis to switch between chemoattraction and chemorepulsion

**DOI:** 10.1101/2025.10.28.685129

**Authors:** Grace K. Luettgen, Brian A. Camley

## Abstract

White blood cells can be guided to targets by chemoattractant signals, but this response is complicated, including guidance toward and away from inflammation sites. We model how cells can switch between being attracted and repelled by a chemical signal. We study experiments on malignant B cell lines, which find that depending on their environment, B cells can either be attracted or repelled by signals like CCL19. The presence of chemorepulsion is also dependent on whether the receptors for CCL19 can be internalized via endocytosis. We develop a stochastic model of receptor-ligand binding and internalization where bound receptors drive a nonlinear feed-forward loop of intracellular signaling molecules, which determine the cell’s direction. We recapitulate key experimental results: changing CCL19 concentration or inhibiting receptor internalization can switch the cell’s direction. Our model implies that cells can navigate toward a target concentration of a signal, regulating that target by receptor internalization. We propose experiments to test this idea.

**Statement of Significance:** Immune cells can be either repelled or attracted by certain proteins. This behavior may allow a single protein to both guide cells to injuries and then encourage them to leave. We provide a theoretical model that can explain experiments showing both repulsion and attraction to a single cue. This model also shows how cells can regulate when they switch between attraction and repulsion by changing the rate of receptor proteins on the surface of the cell being brought into the cell.

## I. INTRODUCTION

Chemotaxis, the migration of cells in response to gradients of chemical cues, plays an important role in wound healing, tissue development, and cancer metastasis [1–3]. For example, neutrophils follow chemical signals left by bacteria to the infection site [4]. Chemical guidance cues for lymphocytes can be attractive [5] or repulsive [6], and lymphocytes must resolve multiple competing chemical cues (e.g. endogenous vs infection-induced cues) to successfully navigate to a target [4, 7, 8]. Cells may also sculpt the chemical environment by secretion and degradation of attractants [9–14]. Combinations of chemorepulsive and chemoattractive cues may work together to help localize a moving source [15]. It is also possible for a single ligand to be both chemoattractive and chemorepulsive, depending on the steepness and/or background concentration of the ligand gradient [6, 16, 17]. This switch may be relevant to the observation of lymphocytes developing “reverse migration” away from sites of inflammation [18].

While chemotaxis has been extensively modeled and quantified in both immune cells [19–21] and other eukaryotic cells [22–27], these models essentially all describe chemoattracted cells and the switch between chemorepulsion and chemoattraction is not well understood. Here, we focus on Malet-Engra et al.’s experimental measurements of single JVM3 cells, a type of malignant lymphocyte that can switch between attraction and repulsion to gradients of the protein CCL19 [17]. Malet-Engra et al. tracked single JVM3 cells in a microfluidic device with a 1 mm long track that established a linear concentration gradient. (The main focus of [17] was the emergent chemotaxis of clusters of JVM3 cells, which differs from single-cell behavior; we will only discuss single cells.) When the CCL19 gradient spanned 0 to 25 ng/mL, single cells had only random migration. However, in the 0 to 100 ng/mL CCL19 gradient, single cells persistently moved towards higher CCL19 concentrations (chemoattraction), and persistently moved to lower CCL19 concentrations (chemorepulsion) in the 0 to 500 ng/mL CCL19 gradient. This chemorepulsion appears to be dependent on the ability of the receptor for CCL19, CCR7, to internalize via endocytosis. When CCR7 internalization was inhibited by knockdown of clathrin or dynamin or treatment with dynasore, cells in the 0 to 500 ng/mL CCL19 gradient flipped their direction and were persistently chemoattracted. Malet-Engra et al. also argue that it is the chemotactic gradient strength that leads to reversal, and not the baseline chemoattractant concentration, because cells have random motion in what we call a “high baseline” condition ranging from 400 to 500 ng/mL instead of chemorepulsion as in the 0 to 500 ng/mL case. These results and the concentration ranges in nM are summarized in Table I. While much work has been done to model these experiments and related questions of how groups of cells can cooperate to follow chemical gradients, potentially overcoming chemorepulsion [28–33} the origins of the switch between chemoattraction and chemorepulsion and the relevance of internalization is not well understood, and these questions have recently been highlighted as “still-unexplained” [34].

**TABLE I.** Summary of key experimental results of [17] for single cells. We have converted to nM using 10.993 kg/mol as the molecular weight of CCL19 (primary accession number: Q99731) [35].

| Concentration range (ng/mL) | Concentration range (nM) | Internalization? | Direction |
| --- | --- | --- | --- |
| 0–25 | 0–2.3 | Yes | Neither |
| 0–100 | 0–9.1 | Yes | Chemoattracted |
| 0–500 | 0–45.5 | Yes | Chemorepulsed |
| 0–500 | 0–45.5 | No | Chemoattracted |
| 400–500 | 36.4–45.5 | Yes | Neither |

How can the same chemical cue lead to opposing behaviors and what role does receptor internalization play? We extend a model of bidirectional chemotaxis of growth cones [16] to understand the experiments of [17]. We assume that bound CCR7 receptors drive a nonlinear feedforward loop and show that this model can replicate the internalization-dependent bidirectional migration of JVM3 cells [17] summarized in Table I. In Section II A, we describe our stochastic model for a CCR7 receptor that can bind ligand, unbind ligand, and internalize. Bound receptor acts as the signal for a nonlinear feed-forward loop (Section II C); the nonlinearity allows the cell to switch directions depending on signal strength. Inhibiting internalization increases the signal, which can switch the cell’s attraction (Section II D). Using a shallow gradient approximation, we propagate the noise from the stochastic receptor dynamics to the cell’s final motion and compute the signal-to-noise ratio for cell migration (Section II E), predicting how accurately cells will chemotax in different chemical gradients. Finally, in Section II F, we predict the existence of stable and unstable fixed points in taxis and show how additional sources of noise may affect our model in Section II G. Importantly, our model can reproduce chemoattraction and chemorepulsion for single cells, and does not require the presence of self-generated gradients or multiple signals [7, 11, 14]. This model also demonstrates a role for receptor internalization in regulating concentration-dependent migration patterns. Our work extends [16], which assumes that a deterministic ligand concentration is the direct input to the nonlinear feedforward loop, by treating the input to the loop as the stochastic number of ligand-bound receptors on the surface of the cell. In making this distinction, we propose that by decreasing the number of receptors on the cell surface, receptor internalization can regulate the effective concentration that a cell senses and more precisely tune the ligand concentrations at which a cell is chemoattracted or chemorepulsed.

## II. MODEL AND RESULTS

### A. CCR7 receptors switch between unbound, bound, and internalized states

Because internalization of CCR7 receptors plays a critical role in the bidirectional chemotaxis of JVM3 cells, we begin by developing a simple model of internalization. CCL19 is known to induce internalization of CCR7. Upon internalization, the CCL19 ligand is degraded inside the cell, while CCR7 is recycled unbound to the surface [36–38]. CCL19 is essential to internalization – in the absence of CCL19, little to no CCR7 internalizes [37, 38]. The simplest way to incorporate these results is to assume that only ligand-bound receptors internalize, and that they are recycled to the surface as unbound receptors (Fig. 1A). This leads to a model with four key rates: 1) unbound receptors in a ligand concentration [L] bind ligand at rate *k*_on_[L], 2) bound receptors unbind ligand with a rate *k*_off_, 3) bound receptors internalize with a rate *k*_in_, and 4) internalized receptors recycle to the surface unbound with a rate *k*_rec_. We will often also work with *K*_D_ = *k*_off_*/k*_on_ – the dissociation constant of the receptor-ligand binding. The master equation for the probability of a single receptor’s state is then:

**FIG. 1.**
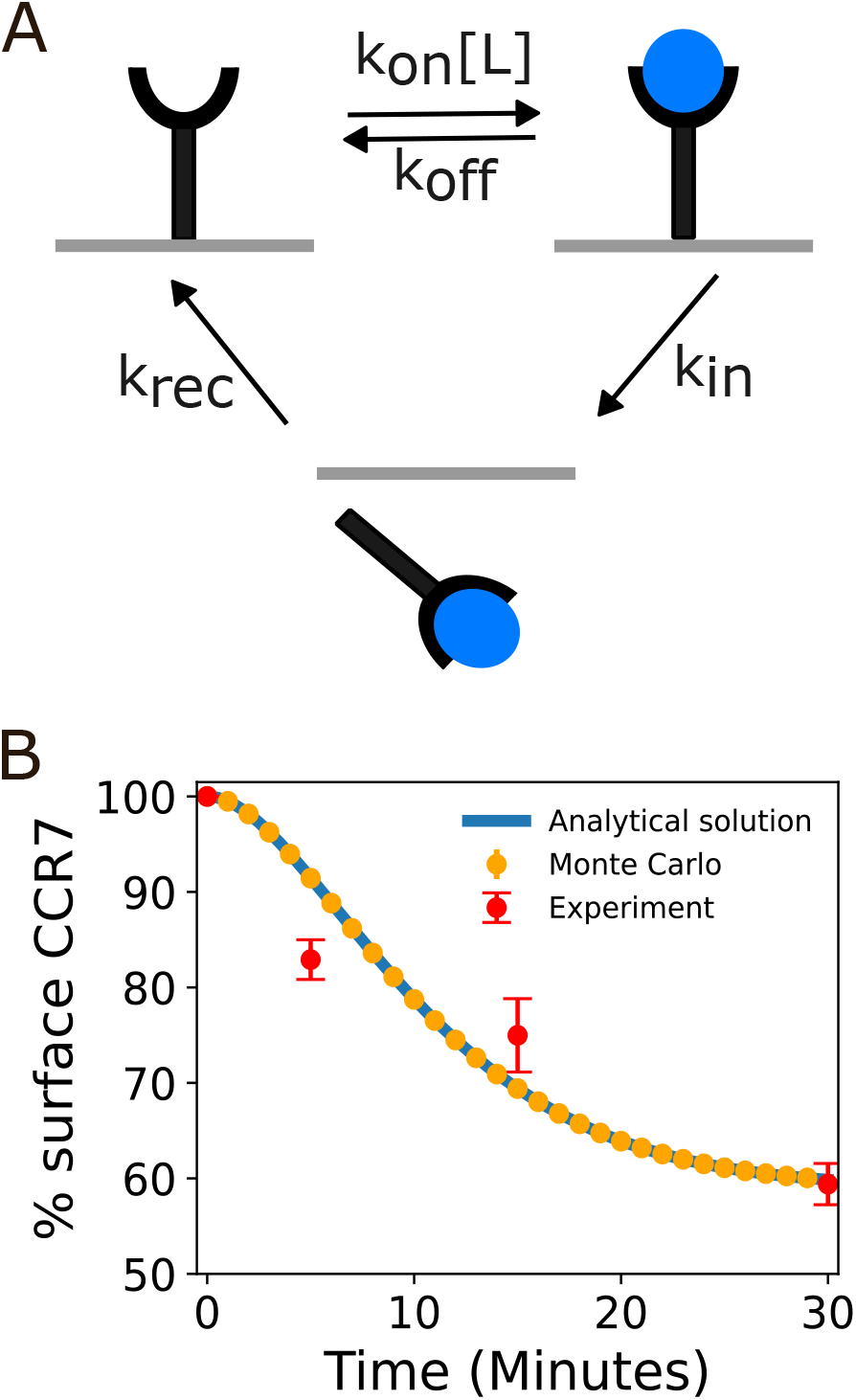
A) Schematic of model of receptor states and transitions of CCR7 receptors (black) and the CCL19 ligand (blue). B) Percent of CCR7 receptors remaining on the surface of a cell exposed to 200 nM of CCL19 over 30 minutes. The blue line is the analytical solution 100 × *p*_s_, where *p*_s_ = *p*_u_ + *p*_b_ is the probability of the receptor being on the surface, the orange points are the average from 100 Monte Carlo simulations, and the red points are experimental data from [17]. Parameters used are given in Table II.

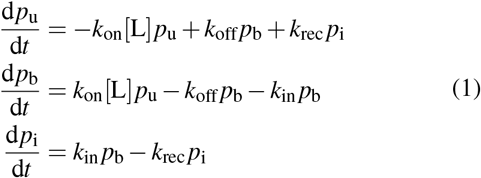

where *p*_u_ is the probability that a receptor is unbound, *p*_b_ is the probability that a receptor is bound, and *p*_i_ is the probability that a receptor is internalized. Solving these equations at steady-state and using *p*_u_ + *p*_b_ + *p*_i_ = 1 we find

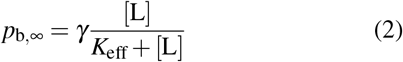

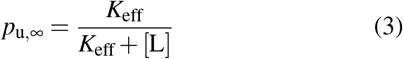

where 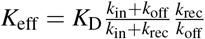 is the effective dissociation constant and 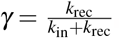. We see that even at saturatingly large ligand concentration, the bound probability is not 100% – *p*_b_ reaches a maximum of *γ*. In the limit of zero internalization, these will converge on the simple ligand-receptor binding fractions:

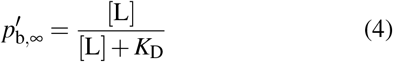

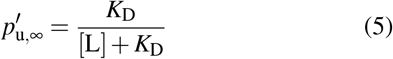

We can also solve Eq. (1) to determine the time-dependent solutions for *p*_u_ and *p*_b_ (Appendix A). For our default parameter values at a wide range of ligand concentrations, the total receptor on the surface (whether bound or unbound) *p*_*s*_ = *p*_*u*_ + *p*_*b*_ can be written as

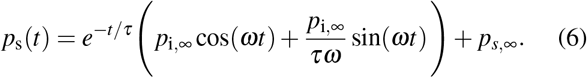

where 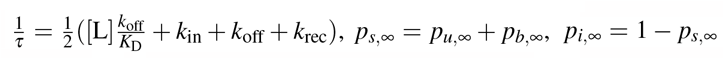 and *ω* is derived in Appendix A. This solution assumes that all receptors are unbound at *t* = 0. We also show in Appendix A that for oscillating solutions in general *ω <* 1*/τ*, and that this constraint makes *τ* the relevant time scale for any choice of parameters in which non-negligible fractions of receptors exist on the surface at equilibrium. In the next subsection, we use Eq. (6) to compare the model to experimental measurements from [17] that measure internalization of CCR7 when cells have been exposed to the CCL19 ligand.

### B. Parameter setting for ligand-receptor model

We set values for *K*_D_, *k*_off_, *k*_in_, and *k*_rec_ using values reported in the literature, and guided by the experimental results on internalization in Fig. 1B. *K*_D_ has been previously reported to be between 1 and 5 nM for CCL19-CCR7 and CCL21-CCR7 complexes, so we use *K*_D_ = 2.5 nM [39–42]. (We note that though [17] use *K*_D_ = 10 nM in their modeling, they choose this value by taking the concentration of maximal migration for pre-B mouse cells in a CCL19 gradient [43], which we find is not necessarily equal to *K*_D_.) For the sake of simplicity, we assume *k*_in_ = *k*_rec_, i.e. *γ* = 1*/*2. This assumption ensures that at sufficiently high concentrations for which 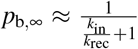 (i.e. [L] ≫ *K*_eff_), half of all receptors are bound and the other half are internalized, a reasonable result given that about 60% of CCR7 receptors remain on the surface of JVM3 cells suspended in 200 nM of CCL19 after 30 minutes [17] (see Fig 1B). We further constrain *k*_in_ and *k*_off_ using chemotaxis data. B cells chemotax most persistently at concentrations on the order of 10 nM [17, 43]. We find later that in our model, cells with functional internalization tend to chemotax most persistently at 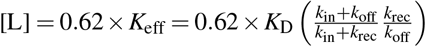 (see Section II E). We thus propose that *k*_off_ is much smaller than *k*_in_ = *k*_rec_. Then 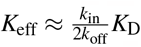, and we find that the ligand concentration that maximizes chemotactic accuracy can be larger than *K*_D_.

We compare the percentage of the receptors on the surface whether bound or unbound, 100 × [*p*_b_(*t*) + *p*_u_(*t*)], for a cell suspended in 200 nM of CCL19 over 30 minutes to the results of [17]. We tuned *k*_in_ and *k*_off_ manually so that they fit experimental data provided by Malet-Engra et al. roughly, as shown in Fig. 1B. We found *k*_in_ = 0.07 min^−1^ and *k*_off_ = 0.002 min^−1^, both of which are within reported ranges for G-protein coupled receptors (GPCRs), the family to which CCR7 belongs [42, 44–46]. Although the value for *k*_off_ is on the lower end for GPCRs, this is expected for a small *K*_D_, and the resulting value for *k*_on_ (0.0008 min^−1^ nM^−1^) is also within the reported ranges for G-protein coupled receptors [45]. The fit in Fig. 1B shows that we have captured the timescale of internalization, though given the relatively limited data, it is certainly possible that more complicated kinetic models could also be supported. We will assume that receptor binding probabilities are at steady state for the remainder of the paper, so we will drop subscripts and write *p*_b,∞_ as *p*_b_, etc. We discuss the validity of this assumption in Appendix C.

### C. Nonlinearity of feedforward loop enables bidirectional chemotaxis

In the experiments of [17], a cell is placed in a linear gradient of CCL19, i.e. ligand concentration is [L] = [L_mid_] + *g* × *y*, where *g* is the gradient slope and [L_mid_] is the ligand concentration at the device midpoint *y* = 0. In our model, the probability of a receptor being bound increases monotonically with ligand concentration (Eq. (2)) – so there are usually more receptors bound on the side of the cell with higher CCL19. For the cell to interpret this signal and decide to travel toward higher or lower ligand concentration, there must be some downstream processing of the signal. We assume that the bound receptor acts as a signal for a nonlinear feedforward loop, regulating a local activator A and inhibitor I, which both regulate a molecule R that leads to the eventual direction of the cell – the cell travels in the direction of higher R (Fig. 2A). This component of the model is a direct adaptation of the approach of [16] to explain bidirectional chemotaxis in growth cones, though we note various feedforward models with differing properties have also been proposed as an element of eukaryotic chemotaxis in other contexts [25, 28, 47–51].

**FIG. 2.**
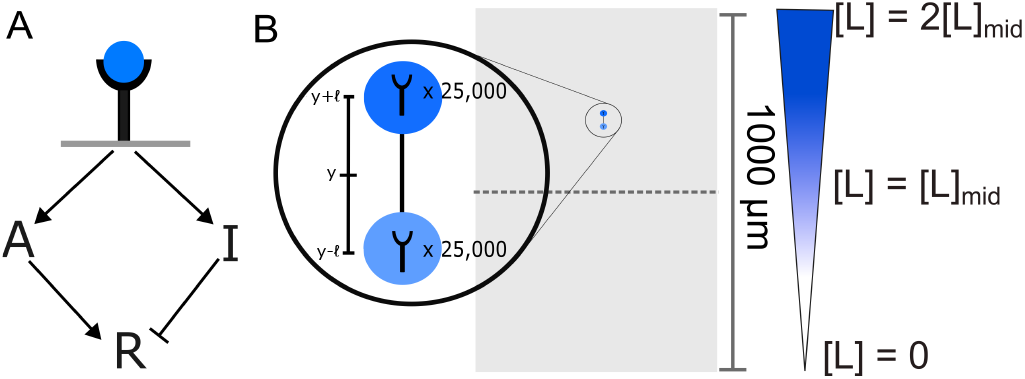
A) A nonlinear feed-forward reaction regulates R – which sets the direction the cell eventually travels – via an activator molecule, A, and an inhibitor molecule, I. B) We use a two-compartment model to approximate the cell such that half of the receptors are located at *y* + *ℓ* and half are located at *y* − *ℓ* relative to the cell’s center at *y*. We assume the cell has 50,000 receptors total. We also illustrate the assumed device geometry ranging from −500 µm to 500 µm (not to scale) with a linear gradient ranging from [L] = 0 to [L] = 2[L_mid_]

Essentially, the idea of the feedforward loop model is that A and I regulate R in opposite ways, and that A and I have different nonlinear dependences on the signal – the number of bound receptors. Then, at different levels of signal, it is possible for A to increase slowly as the number of bound receptors increases, while I increases relatively quickly as a function of the number of bound receptors, leading to R decreasing with an increasing number of bound receptors – and chemorepulsion. To make this more concrete, let’s consider a two-compartment model of the cell similar to that of [52–54], where half the cell’s receptors are located at *y* = +*ℓ* and half are located at *y* = −*ℓ* (Fig. 2B). The ligand concentration [L] is linearly increasing with *y*.

We assume that both A and I are locally regulated by the bound receptors in each compartment and that the local concentrations of A and I have Hill form,

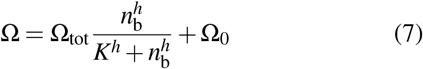

where Ω could be A or I, and where *h* is the Hill exponent, *K* is the half-maximum point and *n*_b_ is the number of receptors bound locally. For our two-compartment approximation, 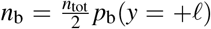 in the front half of the cell, and 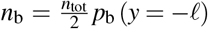 in the back half of the cell. Since *n*_tot_ is just a constant, we can divide *n*_tot_*/*2 out from the numerator and denominator of Eq. (7), effectively absorbing it into *K*. Lastly, we are only concerned with relative comparisons between front and back – so we don’t need to keep track of the overall scaling factor Ω_tot_. This lets us assume that A and I have the form:

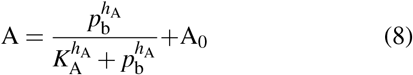

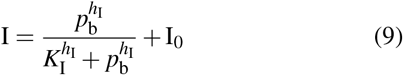

where we have assumed that the A and I reactions can have different Hill exponents *h*_A_ and *h*_I_, half-maximum points *K*_A_ and *K*_I_, and basal levels A_0_ and I_0_. When we later develop stochastic calculations, we will use the actual fraction of receptors bound, i.e. *f*_*b*_ = *n*_b_*/*(*n*_tot_*/*2), in one compartment of the cell instead of *p*_b_.

We assume R is activated by A and inhibited by I [50],

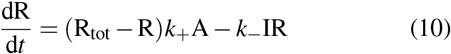

Here R_tot_ represents the total concentration of the response molecule, whereas R represents the concentration of the active response molecule, so R_tot_ = R + R_inactive_. We assume that in the absence of ligand, the steady-state response is completely inactive (R_tot_ = R_inactive_). To enforce this assumption we set A_0_ = 0 and let I_0_ be non-zero, although other parameter choices are certainly possible.

Then, we can write the response R in each compartment at steady state as a function of the probability of a receptor in that compartment being bound:

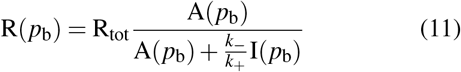

which becomes, after plugging in Eq. 8-9 and setting A_0_ = 0,

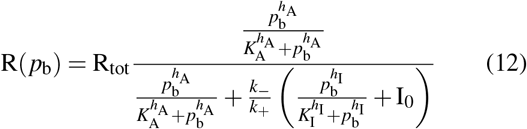

If cells move in the direction of higher R, then we can assume the cell moves in the +*y* direction (higher CCL19) when 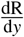 is positive, and the −*y* direction when 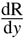 is negative. Because *p*_b_ increases monotonically with *y*, this is equivalent to saying that the cell is chemoattracted if 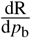 is positive and chemorepelled if 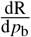 is negative.

The form of Eq. (12) as a ratio of two nonlinear functions can lead to multiple switches between chemorepulsion and attraction. We show an example in Fig. 3A in which I undergoes a rapid transition between its basal level and a high level as *p*_b_ increases (i.e. a large *h*_I_), and A slowly saturates as *p*_b_ increases. We see that at small *p*_b_, I does not change much, while A increases – leading to an increase in 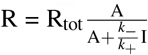. Then, at the transition point where *p*_b_ ≈ *K*_I_, I increases rapidly with *p*_b_, leading to a decrease in R. Then, as I saturates, while A continues to increase, R will increase again. We plot dR/d*p*_b_ for this case in Fig. 3B, seeing that we have chemoattraction at small *p*_b_ and large *p*_b_, but in the intermediate range where dR/d*p*_b_ is negative, there will be chemorepulsion; this chemorepulsive region is shaded in blue throughout Fig. 3. The values of the rate constants we use here are listed in Table II – however, we caution that they are not particularly important. We will later find (Section II E) that the specific rates do not control the signal-to-noise ratio, and the key factor that matters is the sign of dR/d*p*_b_, i.e. the location of the blue chemorepulsive region. To find the zeros of dR/d*p*_b_, we first compute

**FIG. 3.**
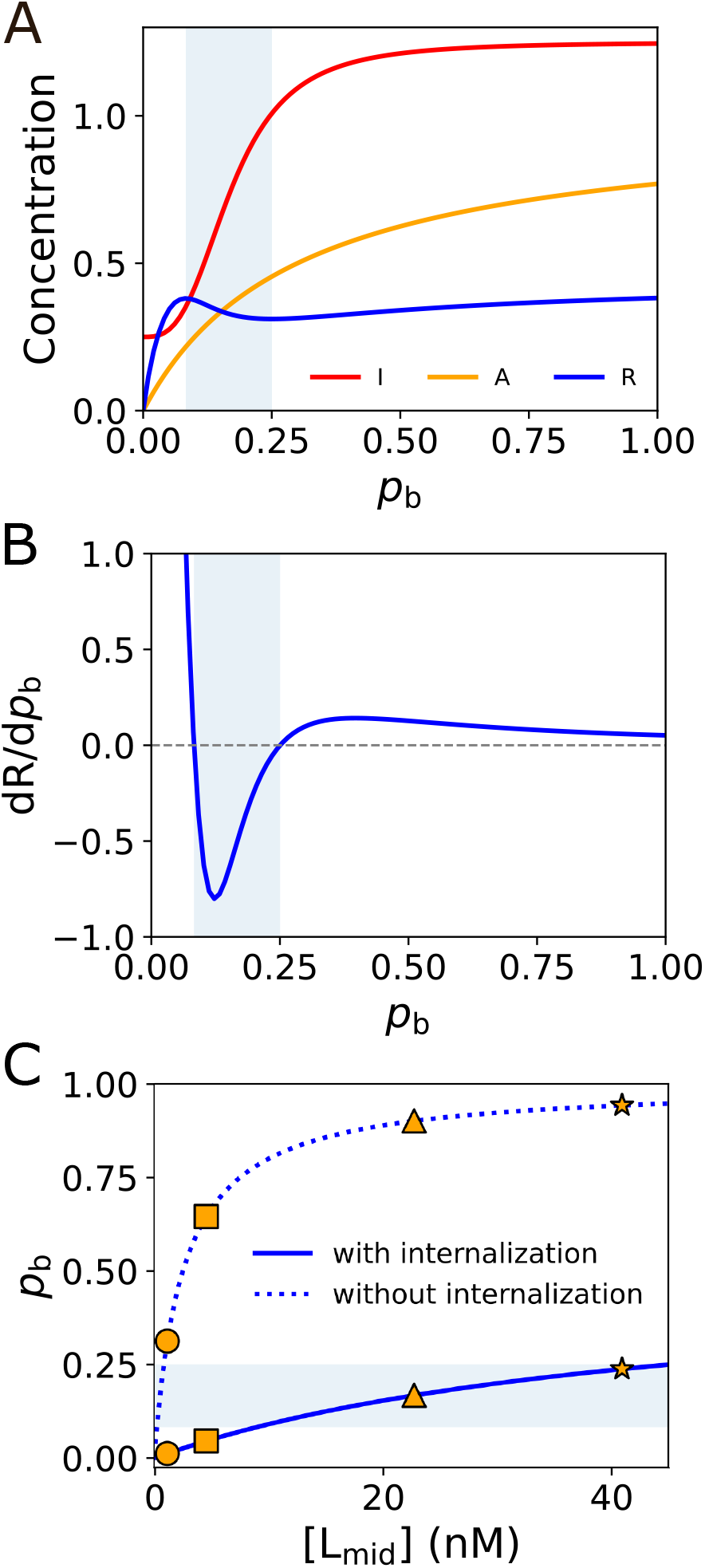
A) A, I, and R as a function of *p*_b_. B) 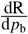 as a function of *p*_b_ shows a region of chemorepulsion (negative 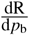). C) Probability a receptor is bound when internalization is (solid line) and is not (dotted line) present. The circle, square, triangle, and star indicate the midpoint concentrations of the 0 to 25 ng/mL, 0 to 100 ng/mL, 0 to 500 ng/mL and 400 to 500 ng/mL gradients respectively. Parameters used are in Table II. Throughout, the blue shaded area indicates the chemorepulsive regime where 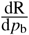 is negative.

**TABLE II.** Default model parameters. Any change from these is explicitly noted.

| Constant | Description | Value |
| --- | --- | --- |
| $K_D$ | Dissociation constant for CCR7-CCL19 binding | 2.5 nM |
| $k_{in}$ | CCR7 internalization rate | $0.07 \text{ min}^{-1}$ |
| $k_{rec}$ | CCR7 recycling rate | $0.07 \text{ min}^{-1}$ |
| $k_{off}$ | Off rate for CCL19 unbinding from CCR7 | $0.002 \text{ min}^{-1}$ |
| $K_A$ | Rescaled Hill concentration for A | 0.3 |
| $K_I$ | Rescaled Hill concentration for I | $0.005^{1/3}$ |
| $I_0$ | Rescaled baseline concentration for I | 0.25 |
| $h_A$ | Hill coefficient for A | 1 |
| $h_I$ | Hill coefficient for I | 3 |
| $\ell$ | Half the length between compartments; $\ell = +2r/\pi$ is the average position of the receptors on half a circular cell of radius $r = 6 \mu\text{m}$ | $3.82 \mu\text{m}$ |
| $n_{tot}$ | Total number of receptors | 50,000 |

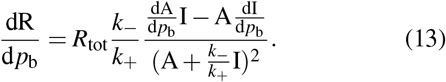

We can see that dR/d*p*_b_ will be zero only if the numerator of the right-hand-side of Eq. 13 is zero, and so these zeros are independent of R_tot_ and 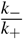. We thus set these parameters to 1 for convenience. In fact, the numerator of Eq. 13 is, up to a positive factor, proportional to 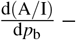 so the zeros of dR/d*p*_b_ are just the zeros of 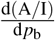. Ref. [16] and other works [28] commonly assume a response R = A/I; we find that the points of switching will be the same whether or not this assumption is made. We note that other choices for *K*_I_, *K*_A_, *h*_I_, *h*_A_, I_0_, and A_0_ can change the number and location of the zeros so that only chemoattraction is possible, or only repulsion, or there is only one switch, etc.; this is explored in depth in [16].

### D. Receptor internalization determines the concentrations at which a cell is chemoattracted/repulsed by modulating the probability a receptor is bound on the cell’s surface

Given our model, how does the concentration of ligand and the presence of internalization change whether a cell is chemoattracted or repelled? We plot the probability of a receptor being bound with and without receptor internalization as a function of ligand concentration in Fig. 3C. With the parameters we have chosen for the feed-forward loop, increasing ligand concentration will bring the probability bound into the blue chemorepelled region at high ligand concentrations if there is internalization, but if there is no internalization, we would predict that at all but very small ligand concentrations (*<* 2 nM or so) cells should be chemoattracted. These results in Fig. 3C show that internalization can switch between chemoattraction and chemorepulsion simply by modulating the number of bound receptors on the surface.

We mark on Fig. 3C the midpoint concentrations corresponding to the 0 to 25 ng/mL, 0 to 100 ng/mL, 0 to 500 ng/mL, and 400 to 500 ng/mL experiments of [17] with a circle, square, triangle, and star respectively. By examining the probability a receptor is bound at these midpoint concentrations, we would predict that cells at the midpoint of the 0 to 500 ng/mL experiment would be attracted in the absence of internalization, but repelled in its presence – exactly as seen in the experiments. However, this model would also predict that you would see chemoattraction at the midpoint of the 0 to 25 ng/mL gradient and chemorepulsion at the midpoint of the 400 to 500 ng/mL gradient where the experiments see no directed motion. We will see below that cells in these latter two experiments may lose directionality because of stochastic noise (Section II E and Section II G).

### E. Binding stochasticity can limit chemotactic accuracy

We have assumed that the cell crawls toward the side that has higher R – i.e. if ΔR = R(*y* = *ℓ*) −R(*y* = −*ℓ*) is positive, it travels in the +*y* direction. However, because there are a finite number of receptors on the cell’s surface, the stochasticity of receptor-ligand binding and receptor internalization means R will fluctuate, so even if our calculation shows that dR/d*p*_b_ is positive, ΔR may often be negative due to these fluctuations. We characterize the extent of these fluctuations by the signal-to-noise ratio, here as the mean ΔR over its standard deviation,

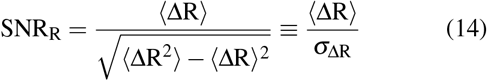

When the magnitude of SNR_R_ is much smaller than 1, we expect the noise to overwhelm the cell’s ability to follow the gradient. We note that Eq. (14) is defined so that it is *signed* – a negative SNR_R_ shows the presence of chemorepulsion.

We know that the signal that drives R is determined from the ligand-receptor binding interactions. We propagate uncertainty to find SNR_R_ in terms of the statistics of ligand binding. We can write 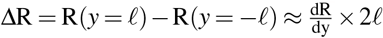. The standard deviation of ΔR can be written as 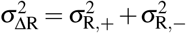, where 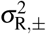 are the standard deviations of R in the compartments at *y* = ±*ℓ*, and we treat the compartments as independent. We can propagate error from the fluctuations in the number of receptors bound in the two compartments as 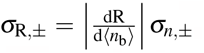, where *σ*_*n,±*_ are the standard deviations of the number bound in each compartment. The SNR approach will be most useful when the noise from the stochastic binding is limiting to the cell’s accuracy – i.e. when the difference between front and back is small. In this shallow gradient limit, we work to smallest order in the difference between front and back, i.e. we assume the differences in the variances between the two compartments can be neglected, 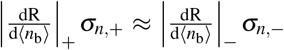. We then find 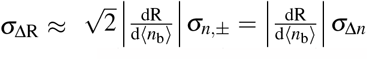, where the derivatives are taken at the cell midpoint and we implicitly defined *σ*_Δ*n*_ as the standard deviation of Δ*n*_b_ = *n*_b,+_ − *n*_b,−_.

Given 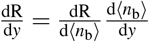, we see:

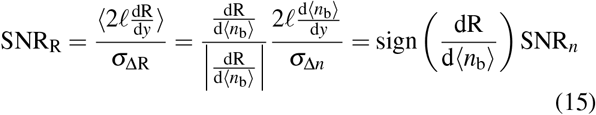

where 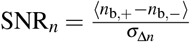 is the equivalent signal-to-noise ratio in the shallow gradient limit for the bound receptor. In other words, the transformation from the number of receptors bound to R only changes the sign, and not the magnitude of the signal-to-noise ratio. This calculation, we note, assumes that the feedforward loop does not add additional noise.

To determine SNR_*n*_, we note that, as we have assumed the receptors are acting independently, the number of bound receptors in a compartment with *n* receptors will follow a binomial distribution, and will thus have mean *np*_b_ and standard deviation 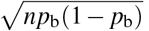. The mean difference between *y* = ±*ℓ* will then be 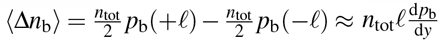, and the variance 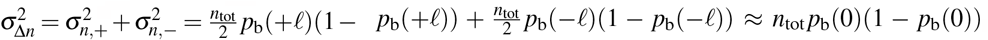, where the last approximation assumes a shallow gradient. Given the mean and standard deviation of Δ*n*_b_, which we can work out from Eq. (2), assuming that the receptors are in their steady-state distribution, we can construct SNR_*n*_ as

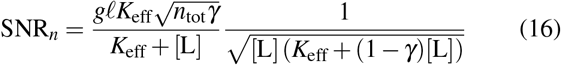

where [L] is the ligand concentration at the cell’s center. SNR_R_ can then be computed as sign 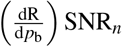. We test that the shallow gradient assumptions and the propagation-of-error calculation are appropriate for our case in Appendix B; we find that a more detailed calculation will smooth some sudden transitions but not change anything essential in our theory. We test the accuracy of the steady-state assumption in Appendix C, and find that (with some important exceptions, and with some caveats on how experiments measuring SNR would have to be done), that the steady-state assumption gives qualitatively correct answers that become quantitatively correct as cell speeds decrease.

We have assumed the chemoattractant profile is [L] = [L_mid_] + *gy*, so if the concentration ranges from 0 to 2[L_mid_] over the channel length of 1000 µm, we find *g* = [L_mid_]*/*500 µm. Given this value of *g*, we find that the numerical maximum of Eq. (16) with *γ* = 1*/*2 (as in our parameters) is at [L_mid_] = 0.62*K*_eff_, i.e. the maximum absolute accuracy occurs a little below the effective dissociation constant *K*_eff_. For our chosen parameters, 0.62 × *K*_eff_ ≈ 28 nM.

For a non-internalizing cell, where *γ* = 1 and *K*_eff_ → *K*_D_,

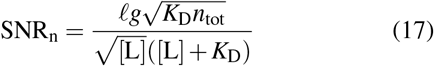

While our signal-to-noise ratio is a relatively simple measure of chemotactic accuracy, we expect, based on the results of [55], that it will be closely related to the Fisher information that is also often used to measure gradient sensing accuracy [23, 56–58].

The sign of SNR_R_ sets the direction in which cells move on average and the magnitude gives a sense for variability about this average. What does our model predict for the experiments of Malet-Engra et al. [17]? Recall, CCR7-internalizing cells in the 0 to 25 ng/mL and 400 to 500 ng/mL gradients moved randomly; CCR7-internalizing cells in the 0 to 100 ng/mL gradient were chemoattracted, CCR7-internalizing cells in the 0 to 500 ng/mL gradient were chemorepulsed, and non-internalizing cells in the 0 to 500 ng/mL gradient were chemoattracted. Fig. 4A shows SNR_R_ for the case where the cell is able to internalize receptors, and Fig. 4B shows SNR_R_ for the case where the cell is not able to internalize receptors. The solid blue lines are the analytical SNR_R_, Eq. (15)-(16), for a cell at the midpoint of the channel as a function of the ligand midpoint concentration [L_mid_], under the assumption that the linear gradient ranges from [L] = 0 nM at *y* = −500 µm to [L] = 2[L_mid_] nM at *y* = +500 µm, i.e. 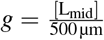. The circle, square, and triangle indicate the SNR_R_ at the midpoint concentration for the 0 to 25 ng/mL, 0 to 100 ng/mL, and 0 to 500 ng/mL gradient cases, respectively. We see solid qualitative agreement with [17]. The 0 to 25 ng/mL case has a low |SNR_R_| relative to the 0 to 100 ng/mL and 0 to 500 ng/mL cases, indicating that cells at the center of the 0 to 25 ng/mL gradient are less directional than cells at the center of the 0 to 100 ng/mL or 0 to 500 ng/mL cases. This prediction aligns well with the experimental result that cells in the 0 to 25 ng/mL gradient move randomly. Cells at the center of the 0 to 100 ng/mL gradient have a large, positive SNR_R_, indicating that cells in the 0 to 100 ng/mL gradient will be accurately chemoattracted, and cells at the center of the 0 to 500 ng/mL gradient have a large, negative SNR_R_, indicating that cells in the 0 to 500 ng/mL gradient will be clearly chemorepulsed. Moreover, non-internalizing cells at the center of the 0 to 500 ng/mL gradient have a large, positive SNR_R_, indicating that when receptor internalization is inhibited, JVM3 cells are chemoattracted with high accuracy in the 0 to 500 ng/mL case.

**FIG. 4.**
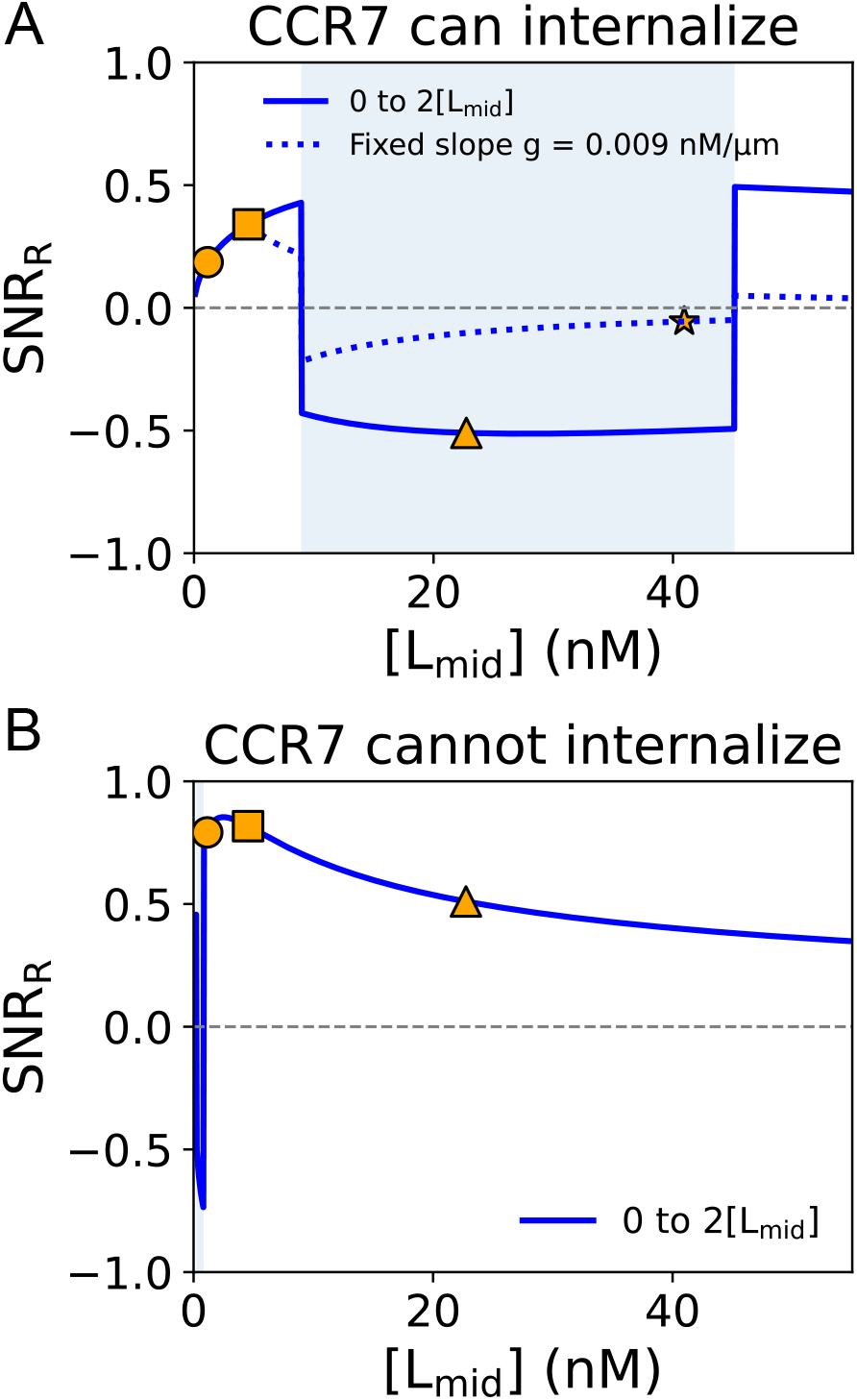
A) SNR_R_ when receptors can internalize. The solid line is SNR_R_ of a cell at the center of the track when the concentration changes linearly from 0 to 2[L_mid_] over the length of the 1 mm track. The dotted line indicates the SNR_R_ of a cell at the center of the track with a gradient steepness of 0.0091 nM/µm and a midpoint concentration given by [L_mid_]. The triangle, square, and circle indicate the SNR_R_ at the midpoint concentration for the 0 to 500, 0 to 100, and 0 to 25 ng/mL concentration gradients. The star indicates the SNR_R_ at the midpoint concentration for the 400 to 500 ng/mL experiment. Note that the dotted line does not span to the smallest range of [L_mid_] as this would have made the concentration negative at one end of the device. Parameters are the same as in Table II. B) The analytical SNR_R_ for the non-internalizing case. For both plots, the blue shaded region indicates the chemorepulsive regime and the dashed gray line is SNR_R_ = 0.

The dotted blue line in Fig. 4A is the analytical result for SNR_R_ in the “high baseline” case, when the device has a fixed gradient of *g* = 0.0091 nM/micrometer, which is the steepness of the 400 to 500 ng/mL experiment. The star shows the midpoint concentration of the 400 to 500 ng/mL gradient experiment. As expected, this SNR_R_ is low relative to the 0 to 500 ng/mL and 0 to 100 ng/mL cases, indicating that cells in the 400 to 500 ng/mL case move less directionally. The low directionality of the high baseline case reflects the relatively small percentage change in signal between the back and the front in the 400 to 500 ng/mL experiment, and the high noise at this concentration range.

Our analytical model replicates the key experimental results for cells at the center of the device in the five conditions in Table I: CCR7-internalizing cells are chemoattracted in the 0 to 100 ng/mL gradient, CCR7-internalizing cells are chemorepulsed in the 0 to 500 ng/mL gradient, non-internalizing cells are chemoattracted in the 0 to 500 ng/mL gradient, and cells move less directionally in the 0 to 25 and 400 to 500 ng/mL gradients. These results show our model is sufficient to explain this switching behavior, at least at the qualitative level.

One potential concern is that we do not see a large quantitative difference in SNR between the 0 to 25 ng/mL condition and 0 to 100 ng/mL condition (circle and square in Fig. 4), while experiments see no measurable chemotaxis for the smaller concentration. We expect that this is because our model neglects other sources of noise – including an additional generic source of noise strongly suppresses chemotaxis in the 0 to 25 ng/mL experiment (Section II G below).

### F. Model parameters predict stable and unstable equilibrium points

In Fig. 4, we are summarizing the signal-to-noise of an experimental condition by its value at the midpoint of the track. However, because the concentration changes along the track, a cell’s SNR will change along the track, including possibly changing sign. Initially, this seems like a relatively minor issue. If we randomly distribute cells around the track, essentially all CCR7-internalizing cells in the 0 to 100 ng/mL gradient and non-internalizing cells in the 0 to 500 ng/mL gradient are chemoattracted for our chosen parameters. Additionally, 80% of CCR7-internalizing cells in the 0 to 500 ng/mL gradient should be chemorepulsed. However, the change of sign of attraction along the track may have potentially interesting consequences. Our model assumes that the probability of a receptor being bound *p*_b_ determines whether a cell is chemoattracted or repelled, and that *p*_b_ depends on [L]. Holding other parameters fixed, there must be some equilibrium [L] for which the cell is neither chemoattracted nor chemorepulsed. This feature would be a necessary part of any concentration-dependent model of bidirectional migration. These equilibrium points are at the concentrations where SNR_R_ changes sign in Fig. 4A – roughly 9 nM and 45 nM. For cells that are able to internalize receptors, the equilibrium at 9 nM is stable. Cells starting at concentrations less than 9 nM will tend to move towards increasing concentrations (SNR_R_ *>* 0), and cells starting at concentrations greater than 9 Nm will tend to move towards decreasing concentrations (SNR_R_ *<* 0). The equilibrium at 45 nM can similarly be seen to be unstable – cells starting at ligand concentrations below 45 nM move toward lower concentrations, and cells starting at ligand concentrations above 45 nM move toward higher concentrations.

The equilibrium points of 9 nM and 45 nM are not strongly constrained by the experimental data – the chemorepulsion switch could clearly move in Fig. 4A while still being consistent with the experimentally-determined concentration dependence of the migration direction. We can at most say that it is likely a stable equilibrium in our model that is consistent with experiment should be in roughly the 4.6 nM to 23 nM concentration range.

If we plot SNR_R_ as a function of the cell’s position, we predict that there is actually a stable point near one end of the device used by [17] in their 0 to 500 ng/mL experiment (Fig. 5). Is this consistent with the experiment? For our model parameters, CCR7-internalizing cells in the 0 to 500 ng/mL gradient should be chemorepulsed if they start migrating at *y >* −303 and *y <* 492 micrometers, meaning that CCR7-internalizing cells randomly distributed over the 0 to 500 ng/mL gradient should have a negative SNR_R_ about 80% of the time. This prediction aligns well with the experimental result that 84% of cells in the 0 to 500 ng/mL gradient were chemorepulsed [17]. However, many cells while being chemorepulsed migrated the full length of the track in the 0 to 500 ng/mL experiment, including ~200 micrometers of what we predict to be the chemoattractive region. Moreover, there is no evidence that cells reach an equilibrium near the low-concentration end of the track in the 0 to 500 ng/mL gradient. The presence of a chemoattractive region in this experiment is not just a feature of our specific parameter choices but is often true – if the ligand concentration [L] alone determines cell migration direction, then the 0 to 500 ng/mL gradient for CCR7-internalizing cells should contain both chemoattractive and chemorepulsive regions, since a 0 to 100 ng/mL region is contained within it. We do not see immediate evidence of cells undergoing location-dependent chemoattraction and chemorepulsion in the experimental movies of [17]. However, we emphasize that it is not trivial what the behavior of a cell near an attractive equilibrium point will be. First, it takes time for receptors to reach equilibrium in a new ligand concentration, as in Fig. 1B. Second, cells can have a memory of a previous chemical cue which can potentially override the new signal [48, 49, 59–61]– the details of which will depend on complicated factors including how the memory couples with the eventual response R. (Cells in tight confinement like in [60, 61] may also be less likely to respond to new signals, though that is not relevant in [17].) Cell speed and polarity may also evolve over an experiment [62]. Depending on these timescales, we might expect oscillations about the attractive equilibrium point, but with different levels of noise depending on the SNR and the memory. The presence of these attractive points might also be masked if they are close to the edge of a device, as in Fig. 5. In these cases if we think of the memory of a cell as playing a role of an effective inertia, the cell may keep going past the attractive point and exit the gradient region of the device. This absence of a clear equilibrium point may suggest that our model is incomplete, or it may be possible to detect the presence of an attractive equilibrium point by further experiments that track the chemotactic velocity of the cells as a function of their initial location, as in the convergent and divergent behaviors found in [7].

**FIG. 5.**
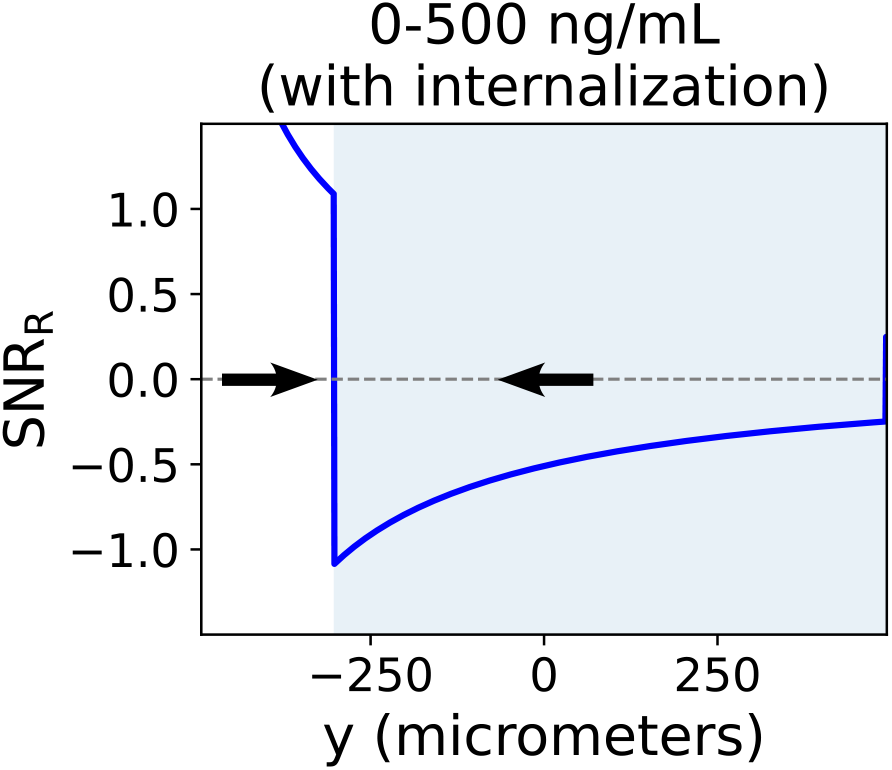
Analytical form for SNR_R_ as a function of the position along a 1mm track showing a predicted stable equilibrium. The blue shaded area indicates the chemorepulsive regime, and the arrows indicate the direction of migration.

Because current experiments only loosely constrain the location of the equilibria, it seems like the best way to design an experiment to observe an equilibrium point would be to try to put the equilibrium near the center of the device. We show some examples of SNR curves as a function of the track length for different gradient conditions and for cells with and without internalization in Fig. 6, where stable and unstable equilibria can be created. However, our concerns noted above about the role of the cell’s memory and polarization also apply in these proposed experiments. One way to discriminate the role of memory in JVM3 cell migration would be to place cells in a chamber such as that described in [48, 63], which allows for rapid control of the gradient. If cells start in a 0 to 500 ng/mL gradient, and then the gradient is suddenly switched to a 0 to 100 ng/mL gradient, one could measure how long the cells are chemorepulsed before switching directions.

**FIG. 6.**
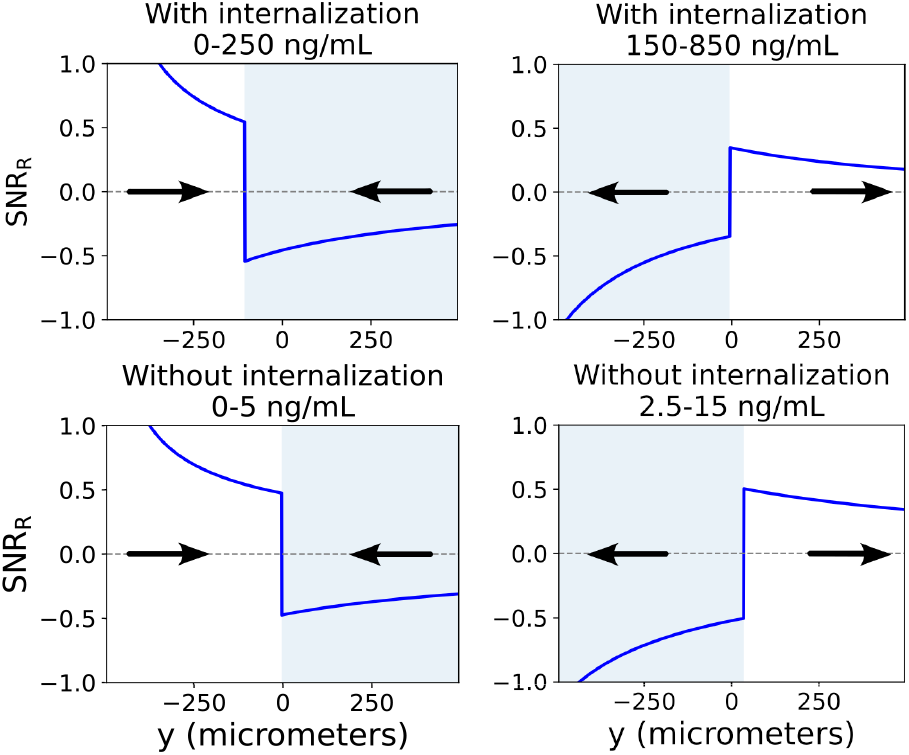
Predictions of chemotactic gradients that would establish stable (left column) and unstable (right column) equilibrium concentrations for CCR7-internalizing cells (top row) and non-internalizing cells (bottom row). The blue curves are the analytical form for SNR_R_ as a function of the position along a 1mm track with a linear gradient of CCL19 that ranges from 0 to 250 ng/mL, 150-850 ng/mL, 0-5 ng/mL, or 2.5-15 ng/mL. The blue shaded area indicates the chemorepulsive regime, and the arrows indicate the direction of migration.

### G. Additional noise sources are highly relevant at low concentrations

We show SNR_R_ as a function of track length for CCR7internalizing cells in 0 to 25 ng/mL, 0 to 100 ng/mL, and 0 to 500 ng/mL gradients and for non-internalizing cells in the 0 to 500 ng/mL gradient – the four primary conditions we have focused on – in Fig. 7 (solid black line). We note that SNR_R_ increases sharply at the end of the track with small *y*. This increase makes sense since at the 0 nM end of the track, the back of the cell is fully unbound and so any signal at the front of the cell allows it to predict the correct gradient direction. Similarly, even just a few binding events in the absence of any background concentration can guide a cell [64]. We do not think that this predicted sharp increase in SNR_R_ at low concentrations is likely to be observed experimentally. If there is any background noise from effector molecules downstream of R, or some additional inhibition of chemotaxis that must be overcome by higher average levels of ligand [65], we would expect the signal to be suppressed or disguised. This SNR at low ligand concentration is also potentially inconsistent with [17], suggesting that cells starting at the low end of the 0 to 25 ng/mL gradient should move persistently towards higher concentrations for some period of time. However, the experiments by Malet-Engra et al. show that JVM3 cell migration is random in the 0 to 25 ng/mL gradient and previous studies have also seen that cells do not chemotax persistently at sufficiently low concentrations [22, 66, 67]. Therefore, it is likely that we are overestimating the SNR_R_ at low concentrations, since we are only taking the noise of ligand-receptor binding and internalization into account. To model this loss of directionality at low concentrations, we can modify our SNRs to include some additional noise, *σ*′, added in quadrature with the noise of ligand-receptor binding:

**FIG. 7.**
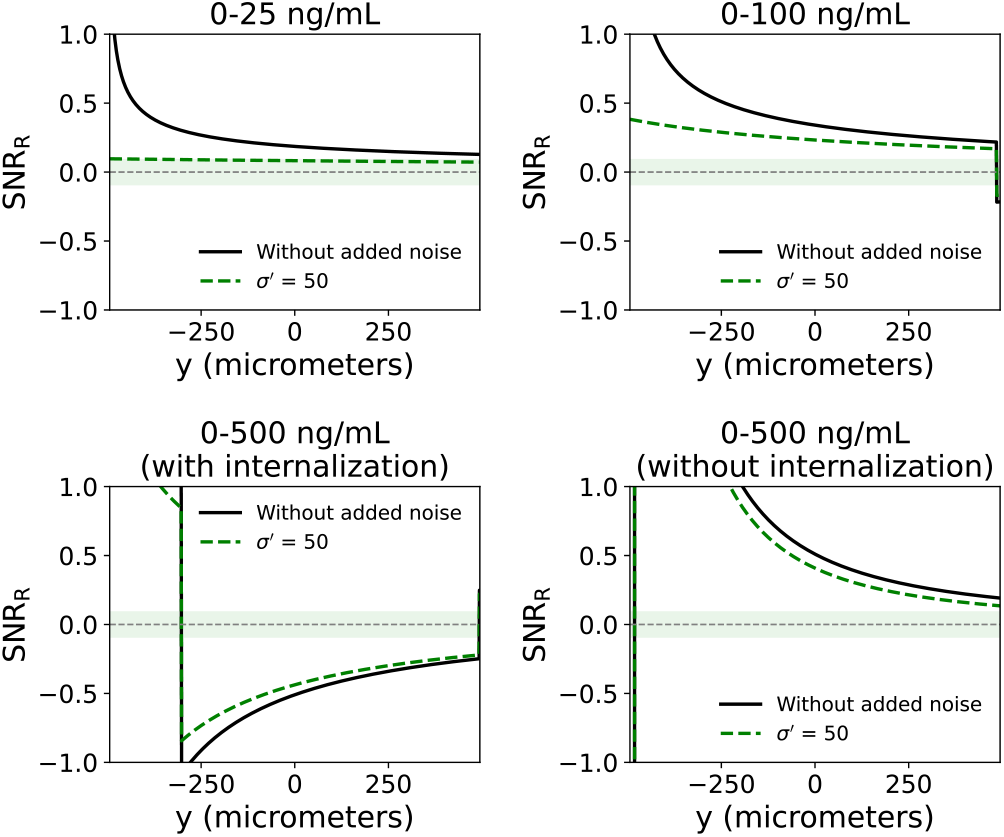
SNR_R_ (solid black) and 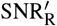 (dashed green, added noise of *σ*′ = 50) of cells at different points along a 1 mm track. The shaded regions are the random migration regimes for the 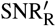 curve–any value of 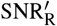 within this shaded region indicates that the cell will move randomly. The maximum absolute value for this shaded region (threshold) is set by the maximum value of 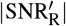 in the 0 to 25 ng/mL gradient. Parameters used are the same as in Table II.

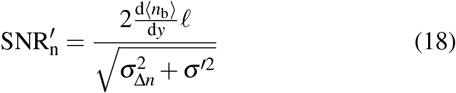

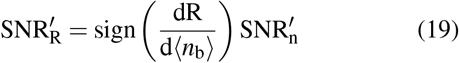

*σ*′ is unitless – we will think of it in comparison to the fluctuations in the number of bound receptors *σ*_Δ*n*_. In Fig. 7 we plot 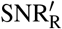 as a dotted green curve; we have chosen *σ*′ = 50. This value is somewhat arbitrary, but we note that the shallowest detectable gradients in Dictyostelium correspond to a front-to-back difference of ⟨Δ*n*_*b*_⟩ ~ 100 bound receptors [67]. We would predict from Eq. 18 that if ⟨Δ*n*_*b*_⟩ ≪ *σ*′, the added noise will suppress chemotaxis independent of any binding noise *σ*_Δ*n*_. In this sense, added noise on the scale of 10-100 is consistent with the idea that cells start to lose the ability to sense gradients of 100 bound receptors. In Fig. 7, we see that the added noise largely affects the two smallest concentration experiments. In particular, in the 0 to 25 ng/mL experiment, the added noise nearly completely suppresses the increase of SNR toward *y* = −500 microns where the concentration goes to zero.

Experiments find only random migration in the 0 to 25 ng/mL experiment, but do find relevant chemotaxis in the 0 to 100 ng/mL experiment. We certainly see in Fig. 7 that the SNR for the 0 to 25 ng/mL experiment is low when the added noise is taken into account. We assume that the highest value of 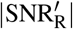 in the 0 to 25 ng/mL gradient is the threshold value for persistent chemotaxis – i.e. any value of 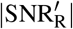 below this threshold indicates random migration and any 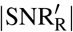 above this threshold indicates at least some observable chemotaxis. We shade the regions where no chemotaxis of any sign should be detected in green in Fig. 7. 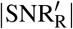 in the 0 to 100 ng/mL and 0 to 500 ng/mL cases is large relative to the threshold value for nearly the entire track as expected, indicating that cells in these gradients will move persistently relative to the 0 to 25 ng/mL case.

Of course, our choice for this threshold is slightly arbitrary – the presence or absence of chemotaxis is not binary, and will depend on additional unmodeled factors, including the timescale over which the cell responds to different concentrations, etc. The main point that we wish to make here is that additional noise ensures both that the SNR does not increase unrealistically as the concentration gets small and that with this additional noise migration is clearly more directional in the 0 to 100 ng/mL and 0 to 500 ng/mL cases than the 0 to 25 ng/mL case.

We also note that the behavior of cells near the low-concentration end of the 0 to 25 ng/mL gradient is also quite sensitive to our assumption that we can treat the receptor dynamics as being at steady state. We see in Appendix C that the SNR in the 0 to 25 ng/mL gradient measured from tracking cells that move at finite speed, will not necessarily have its maximum at low *y* (Fig. 9).

**FIG. 8.**
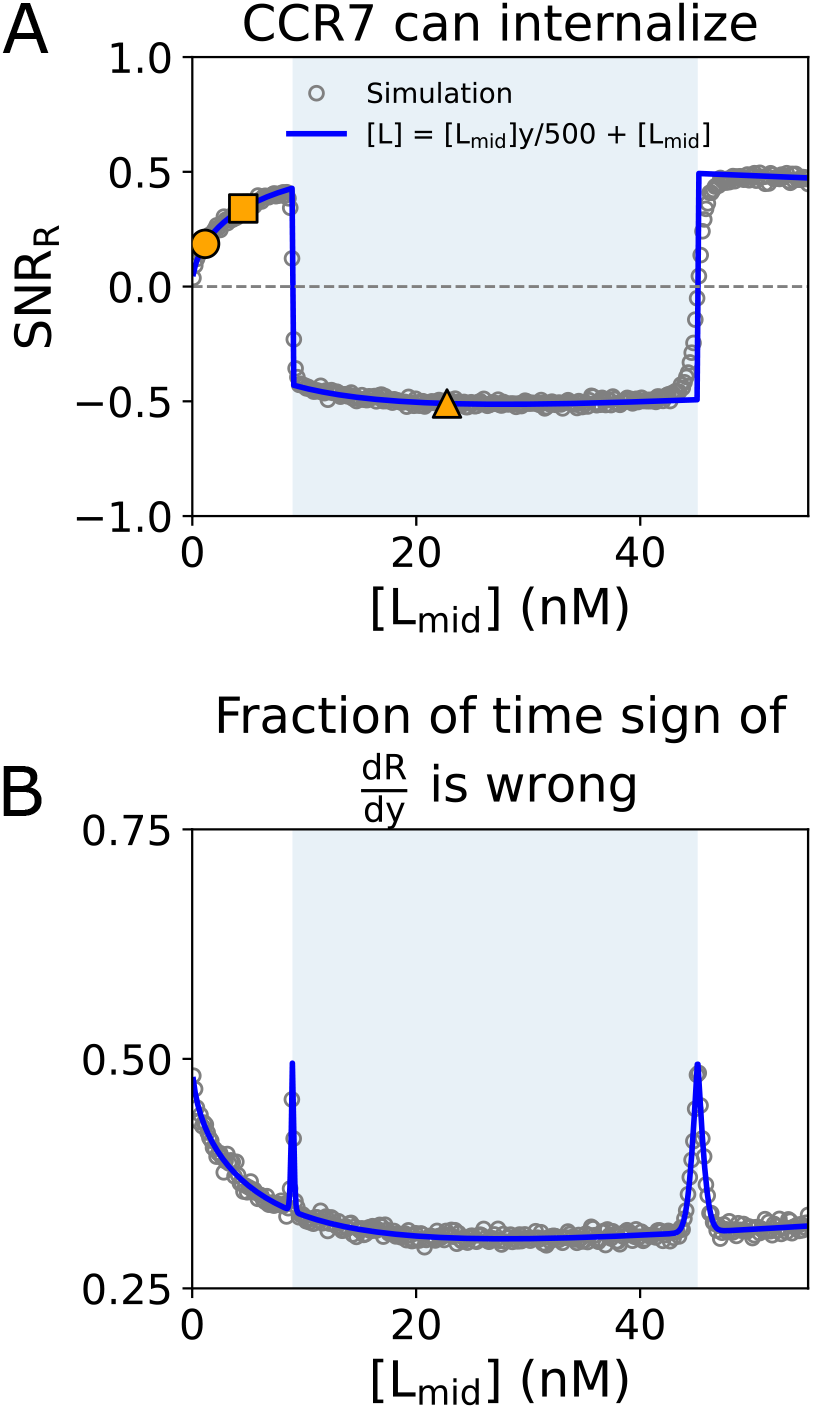
A) Analytical form for SNR_R_ (blue line) overlaid with average of 10,000 simulations (open circles). This numerical result corresponds with the theory result in Fig. 4A. B) Analytical form for the fraction of the time that the cell guesses the sign of 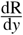 incorrectly (blue curve) overlaid with the average of 10,000 simulations (open circles).

**FIG. 9.**
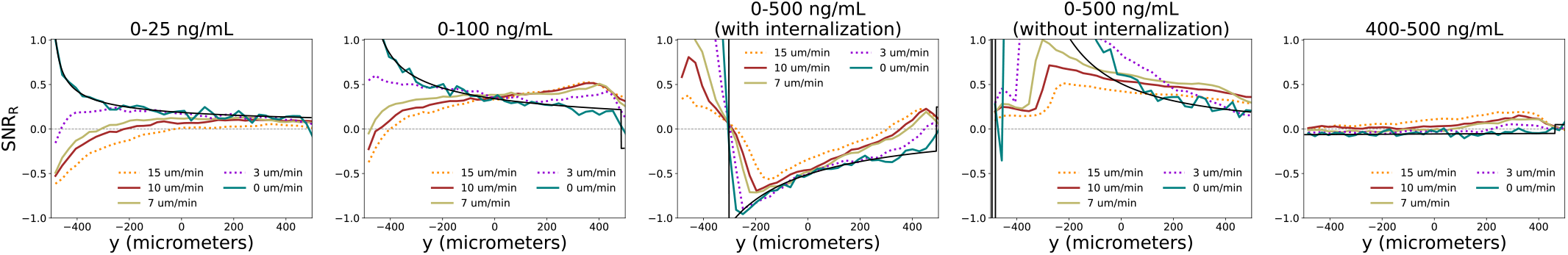
Measurements of SNR_R_. Each plot is constructed from 6,000 stochastic simulation trajectories of single cells (see methods in Appendix C 1). The black line shows the analytical result. Parameters for each simulation are given by Table II.

## III. DISCUSSION

We developed a model of cell migration where the number of bound receptors on the cell surface is the input for a nonlinear feedforward loop that dictates migration direction, which lets the level of receptor internalization modulate the input strength. By calculating the signal-to-noise ratio of the output of the loop, we showed that this model replicates the chemoattraction of CCR7-internalizing cells in response to CCL19 gradients of 0 to 100 ng/mL and the chemoattraction of non-internalizing cells in response to 0 to 500 ng/mL gradients; it also replicates the chemorepulsion of CCR7-internalizing cells in response to 0 to 500 ng/mL gradients, and predicts cells will be less directional in 0 to 25 ng/mL and 400 to 500 ng/mL gradients compared to the 0 to 100 ng/mL and 0 to 500 ng/mL cases. This reproduces – at least qualitatively – the key results of [17]. Our model also predicts that this concentration-dependent switch between chemoattraction and chemorepulsion can potentially lead to stable and unstable equilibrium concentrations that attract or repulse cells. There is no immediate evidence in [17] for these stable points – though our model predicts one might be present toward the end of the device – and so a more detailed comparison between experiment and theory may be needed, including a better understanding of what happens when a cell moves between a chemoattractive and chemorepulsive region.

The assumptions of our model require that CCL19 concentration itself controls the switch between chemoattraction and chemorepulsion. This is contrary to the analysis of [17], who argue that it is not merely high CCL19 concentration that leads to chemorepulsion, because cells are not chemorepelled in the 400 to 500 ng/mL gradient (high baseline concentration, but the same slope of concentration as the 0 to 100 ng/mL experiment). Our interpretation is different – we see that the SNR is small enough for this condition due to the high concentration and low percentage change across the cell that cells are not expected to chemotax at all. Our results, however, do not rule out the idea that gradient strength itself could also create a switch from chemoattraction to chemorepulsion – which could be tested more explicitly by experiments matching SNR but varying gradient strength or concentration.

To make our model’s predictions quantitative, a stronger constraint on certain parameters would be necessary. While we have chosen *k*_in_, *k*_rec_, and *k*_off_ consistent with the experimental measurements of internalization [17] and known values for other G-protein coupled receptors, there is some degree of arbitrariness in these choices. Earlier studies have fit receptor kinetics models to experiments measuring the internalization of CCR7 on T cells as a function of CCL19 concentration [68, 69], but the data in [68] are inconsistent with the timescale of internalization from [17] plotted in Fig. 1B, suggesting cell type dependence on these parameters. We have only focused on the most relevant data from [17] in setting our parameters. More detailed kinetic measurements of CCR7 dynamics on JVM3 cells could challenge our assumptions. We also have neglected within our model other potential roles of endocytosis, including preventing remeasurements of single ligands [70], redistribution of receptors across the cell in response to an applied gradient [71, 72], and increased sensitivity due to cooperativity [73], which could be included in future work. Another implication of our result which could be tested to challenge our model is that other factors which control the number of receptors bound to CCL19 on the surface can induce a switch between chemoattraction and chemorepulsion. For instance, the presence of a competing ligand could take a cell in the 0 to 500 ng/mL experiment out of the chemorepulsive limit by reducing the fraction of receptors bound to CCL19. This mechanism is slightly different from a previously discovered way in which competition between ligands can lead to chemorepulsion [7], which does not require the internal switch from the nonlinear feedforward dynamics we study here. Another possibility to switch chemoattraction and repulsion would be the modification of G protein activity [55], which could alter the signal input to the feedforward loop in our model.

In our signal-to-noise ratio calculations, we assume that the molecular reactions that produce the activator, inhibitor, and response molecules that compose the feedforward loop are noiseless and equilibrate quickly with respect to changes in the number of bound receptors. Accounting for the noise of these molecular reactions would further decrease the magnitude of the signal-to-noise ratio [74– In this case, the magnitude of the SNR might not depend so strongly on ligand concentration, though the switching of the direction of chemotaxis would likely remain robust. While we do not propose intracellular signaling molecules that play the role of activator and inhibitor, we see that within our model that the details of activator and inhibitor matter less than the switching concentrations – the signal-to-noise ratio of the cell is independent of all the details. We argue this can allow the model to have significant predictive value once the switching concentrations are better constrained by experiment. However, to predict the results of specific biochemical interventions, closer connection with detailed pathways [50, 77–81] would be necessary.

Our model and that of [16] assume that the value of A and I at each side of the cell only depend on the local signal (in our case, the bound receptor fraction). This locality contrasts with the well-established local excitation, global inhibition (LEGI) network, which has been proposed to guide neutrophils and Dictyostelium [25, 50, 82], and we could call our approach a “local excitation, local inhibition” (LELI) model. One key motivation of LEGI models is their ability to both describe perfect adaptation to spatially-constant signals while still responding robustly to spatially-graded stimuli [50]. Perfect adaptation, though, is in conflict with the experimental observation that ligand concentration controls the switch between chemoattraction and chemorepulsion in [17] – so perhaps it is not surprising that our model takes a slightly different form. Recent work does suggest the possibility of local inhibitors playing a role in cell polarity and chemotaxis in neutrophil-like cells [83–85], though it has been modeled in a different way than the feedforward approach we use here [84, 85]. It may be possible that different variants of negative feedback contribute to different functions in cell migration, reflecting the complex role of negative signals in chemotaxis [86].

Our model assumes that the CCL19 concentration is established solely by the microfluidic device in [17]. However, it is also possible that the CCL19 concentration along the track may systematically deviate from the linear gradient established by the microfluidic device due to the secretion and/or degradation of CCL19 by JVM3 cells. Some cancer cells and lymphocytes have been known to secrete CCL19, which may affect cell migration [44, 87, 88]. Especially given that the results of [17] depend on internalization of ligand, it is difficult to rule out the possibility that chemorepulsion arises from depletion of CCL19 that somehow reverses the effective gradient that cells see, akin to previous work showing the role of “self-generated” concentration gradients in collective systems [9, 11, 14, 89]. Our model shows, however, that these depletion effects are not necessary to explain chemorepulsion. To discriminate between our hypothesis and one where degradation and secretion play a larger role, it would be necessary to alter the strength of degradation in the experiments of [17]. This could be done by altering the density of cells, or by using microfluidic devices that can quickly replenish chemoattractants, changing media on the timescale of seconds [48, 63, 90]. We note that even if no large-scale self-generated gradients are created in the experiments of [17], the presence of internalization-dependent depletion and secretion could systematically shift the concentrations experienced by the cells so they do not match the simple linear gradient we have assumed; similar effects have been proposed to be able to enhance gradient sensing in *Dictyostelium* [91] and yeast [92]. Concentration changes would change the location of the switches from chemoattraction to chemorepulsion, potentially explaining why no chemoattractive region is apparent in the 0 to 500 ng/mL experiment.

Our results may be more broadly applicable than just to the specific experiments of [17]. Subpopulations of T cells have been shown to chemotax bidirectionally in response to the chemokine SDF-1 [6, 93]. Like JVM3 cells, they are chemoattracted at low ligand concentrations and chemorepulsed at high ligand concentrations. While both attraction and repulsion are mediated by a single receptor type, chemoattraction, but not chemorepulsion, of T cells is inhibited by the tyrosine kinase inhibitors genistein and herbimycin [93]. Within our model, it might be possible to remove chemoattraction by disrupting or saturating the A pathway, ensuring that inhibitory changes upon ligand binding are always larger than attractive ones. Earlier hypotheses about chemorepulsion include a suggestion from Zlatopolskiy et al. that chemorepulsion in T cells could arise from internalized receptors triggering a different signaling pathway from surface receptors [94]. We have shown this may not be necessary, as modulating the number of bound surface receptors via internalization alone enables bidirectional chemotaxis. Krishnan et al. have also proposed various feedforward loops to describe the concentration-dependent bidirectional migration of T cells among other bidirectionally moving cells, but do not link these models to internalization [95]. Our work proposes a testable framework that can link interventions on receptor internalization to the eventual chemotactic behavior across a wide variety of eukaryotic cells.

## AUTHOR CONTRIBUTIONS

GKL developed code and analytical results. GKL and BAC designed research, developed the model, and wrote the manuscript.

## ACKNOWLEDGMENTS

Research reported in this publication was supported by the National Institute of General Medical Sciences of the National Institutes of Health under Award Number R35GM142847. The content is solely the responsibility of the authors and does not necessarily represent the official views of the National Institutes of Health. We thank Wei Wang and Emiliano Perez Ipiña for a close reading of the draft.

## Appendix A Analytical calculation and Monte Carlo simulations of non-equilibrium receptor-ligand dynamics

We use the time-dependent probabilities of a receptor being bound or unbound to confirm that our parameters for *k*_in_, *k*_rec_, *k*_off_, and *K*_D_ are reasonable. Here, we derive the analytical form for the time-dependent probability that a receptor is on the surface (i.e. *p*_s_(*t*) = *p*_u_(*t*) + *p*_b_(*t*)), and we confirm this solution describes the system using Monte Carlo simulations. First, solving Eq. 1, we find that

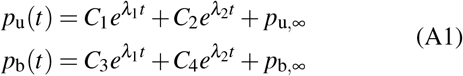

where we can later solve for the constants *C*_1_, *C*_2_, *C*_3_, and *C*_4_ and *λ*_1,2_. The solution for *p*_s_(*t*) is then:

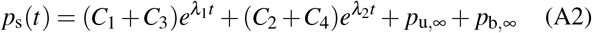

Since we assume all receptors are initially unbound, *p*_b_(0) = 0 and *p*_u_(0) = 1. Substituting these solutions and the steady-state probabilities back into Eq. 1, we find *C*_1_, *C*_2_, *C*_3_, and *C*_4_ are

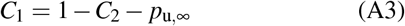

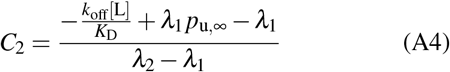

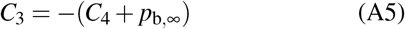

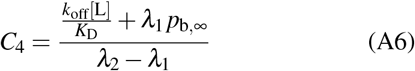

*λ*_1_ and *λ*_2_ can be found by plugging in the form of the solution to the master equation and noting that *p*_i_ = 1 − *p*_u_ − *p*_b_; this shows that *λ*_1_ and *λ*_2_ are the eigenvalues of the matrix:

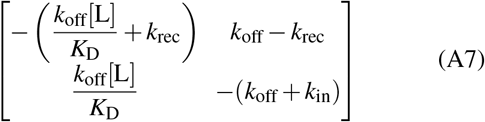

which are, explicitly,

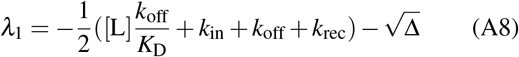

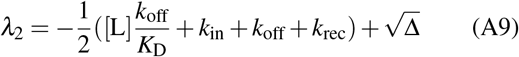

where the discriminant-like term is defined as 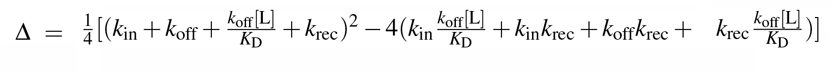. For the parameters we use throughout this paper, Δ is less than zero at some ligand concentrations, causing *λ*_1,2_ to have an imaginary component at those concentrations. However, the imaginary terms in *p*_b_(*t*) and *p*_u_(*t*) cancel, and the solution is real. Can oscillations in receptor occupancy or receptor population on the surface be observed? To investigate this question, we assume that Δ *<* 0 and rewrite Eq. A2 as

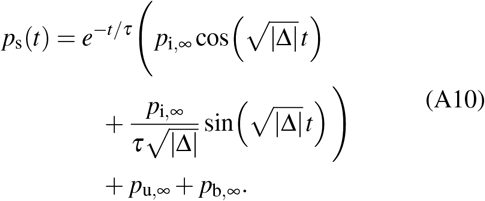

Here we have set 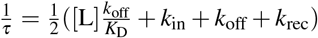. Eq. A10 is Eq. 6 in the main text. The relaxation time, *τ*, is the characteristic timescale that it takes receptor dynamics to reach equilibrium after a perturbation (when Δ *<* 0). Oscillations may become important in nonequilibrium dynamics when the period of oscillation, 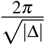, is less than the relaxation time, *τ*. However, the period is always greater than 2*πτ* since 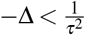 when Δ *<* 0. To see this, note that

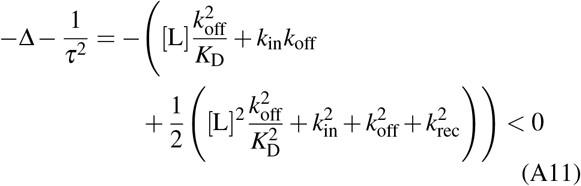

Therefore oscillations are strongly suppressed before completing a full period.

To confirm that our analytical solution describes the system, we also performed Monte Carlo simulations. In these simulations, we modeled the cell as a container with 50,000 receptors in a constant concentration of [L] = 200 nM. All of the receptors are initially unbound and the simulation is allowed to run for 30 minutes in time steps of Δ*t* = 0.01 minutes. Unbound receptors can transform to bound receptors with a probability of *k*_on_[L]Δ*t* at each time step; bound receptors can transform to unbound receptors with a probability of *k*_off_Δ*t* or internalized receptors with a probability of *k*_in_Δ*t* at each time step; internalized receptors can transform to unbound receptors with a probability of *k*_rec_Δ*t* at each time step. We summed the number of bound and unbound receptors at each time step to obtain the number of surface receptors, and plotted this as a percentage of all receptors in Fig. 1B (dotted orange line). The analytical solution and Monte Carlo simulations agree with each other and experimental results, indicating that the analytical solution accurately describes a group of internalizing receptors and that the chosen parameters for *k*_in_, *k*_rec_, *k*_off_, and *K*_D_ are reasonable.

## Appendix B Stochastic simulations confirm analytical results for SNR

In Section II E, we found an analytical formula for the signal-to-noise ratio of the response, SNR_R_, which controls the noisiness of the cell’s ability to be chemoattracted and chemorepelled. However, in deriving it we used a shallow gradient approximation; we also used a propagation-of-error assumption of 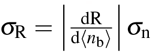, which might be suspect given that our model has rapid changes in R as a function of the number of receptors bound. In this section we test this result using a Monte Carlo simulation of bound receptor statistics in the two-container model of the cell.

Recall, the front container is located at +*ℓ* relative to the cell center and the back container is located at −*ℓ*. Each container contains 25,000 receptors which can be bound, unbound, or internalized. We assume that receptors internalized in one half of the cell are recycled to the surface in the same half of the cell. At equilibrium, the probability that a receptor is bound is simply *p*_b_(*ℓ*) in the front container and *p*_b_ (−*ℓ*) in the back container. For each run of the simulation, we set the 25,000 receptors to be bound with probability *p*_b_(*ℓ*), and similarly set the back container receptors to be bound with probability *p*_b_ (−*ℓ*). Then, we take the fraction of receptors bound in each container and use Eq. (12) to determine R, where the fraction bound is substituted in for *p*_b_. Finally, we take the difference between R in the front and back halves, ΔR. To report SNR_R_, we take 10,000 random samples and then compute the SNR via Eq. (14). The SNR_R_ values at y = 0 for gradients given by 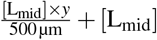 are shown as open circles in Fig. 8A. The solid blue curve in Fig. 8A is the same analytical result for SNR_R_ that was plotted in Fig. 4A. At [L_mid_] far from the transition concentrations (9 nM and 45 nM), the open circles correspond exactly to the analytical result for SNR_R_, indicating that we can obtain SNR_R_ by propagation of error on SNR_diff_.

The SNR_R_’s sign flips at 9 nM and at 45 nM are instantaneous in the analytical curve, but these transitions are smoother in the simulation (Fig. 8A). This smoothness is because the switch in the sign of ΔR is no longer deterministic, but involves small fluctuations in the numbers of bound receptors. In particular, the sign of 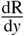 is determined by the difference in bound receptors between the compartments at ±*ℓ, n*_diff_ = *n*_b,+_ − *n*_b,−_, and by the total number of receptors bound in each half, *n*_sum_ = *n*_b,+_ + *n*_b,−_. The total number of bound receptors helps set the sign of 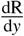 because it determines whether the cell is in the chemoattractive or chemorepulsive regime. For example, a cell in a gradient centered at 10 nM, just into the chemorepulsive regime, may have a larger number of bound receptors at the front than the back, but if the number of bound receptors in both halves happen to be extremely low, then the sign of 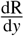 will be positive instead of negative. In other words, the cell guessed the correct gradient direction (*n*_diff_ > 0), but because the sign of ΔR flips at 9 nM and *n*_sum_ is extremely small (*n*_sum_ < *n*_tot_ × *p*_b_(9 nM)), the cell acts as if it is in the low-concentration chemoattractive region of the 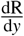 curve, so the cell will be chemoattracted rather than chemorepulsed.

To verify that fluctuations in the total number of bound receptors are responsible for the smooth transitions, we can use *n*_sum_ and *n*_diff_ to calculate the fraction of the time we would expect the sign of 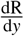 to be incorrect as a function of concentration and gradient steepness. As in Section II E, we assume that *n*_b,+_ and *n*_b,-_ follow binomial distributions with average *µ* and standard deviation *σ*. Because both containers have a large number of receptors, we can approximate these binomial distributions as Gaussians. Then, *n*_diff_ is a Gaussian distribution with average *µ*_diff_ = *µ*_+_ − *µ*_−_ and we can approximate its standard deviation as 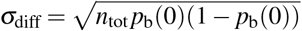 as we did in Section II E. Similarly, *n*_sum_ is a Gaussian distribution with average *µ*_sum_ = *µ*_+_ + *µ*_−_ and we can approximate its standard deviation as 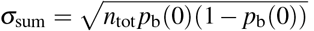. These two variables *n*_diff_ and *n*_sum_, since they are constructed from *n*_±_, are dependent variables with correlation coefficient 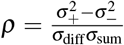 and we can construct a bivariate normal distribution to describe their joint probability:

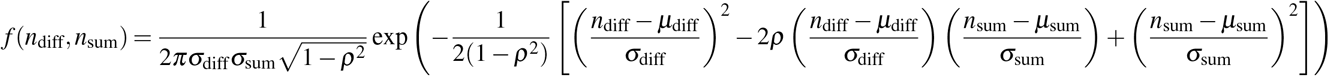

In general, any value of *n*_sum_ less than *n*_tot_ × *p*_b_(9 nM) or greater than *n*_tot_ × *p*_b_(45 nM) places the cell in the chemoattractive regime; otherwise, the cell is in the chemorepulsive regime. As before, any positive value of *n*_diff_ indicates that the cell has guessed the gradient direction correctly. Therefore, for a cell located at the center of a gradient described by 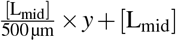 with [L_mid_] *<* 9 nM, the probability the cell guesses the incorrect sign for 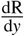 is:

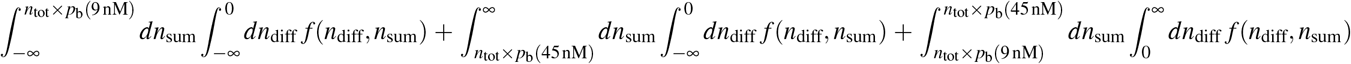

We can perform similar calculations for the chemorepulsive and high-concentration chemoattractive regimes. Fig. 8B plots these analytical calculations for each [L_mid_] (solid blue curve) and compares them to simulation results (open circles). To get the correct sign, the total number of bound receptors just needs to be in the correct regime, and so cells at concentrations far from the transitions have SNR_R_ values that are effectively limited only by gradient sensing noise. However, cells close to the transition concentrations will also be limited by fluctuations in their raw number of bound receptors. We expect the size of this transition region to be about the size *σ*_sum_ of the fluctuations in *n*_sum_ – so the transition from negative to positive at 45 nM is particularly smooth because *σ*_sum_ is higher at higher concentrations. Looking at the fraction of the time the sign of dR/dy is wrong in Fig. 8B, we see two spikes that reach a maximum of 0.5, one at each transition region, with the second being much broader as expected.

We note, in principle, that the front of the cell could have a value of *n*_+_ that puts it in the chemoattractive regime and the back is in the chemorepulsive regime. We have neglected this possibility in computing the theoretical curve, since we are in the shallow gradient limit. We see from the good agreement between theory and simulation in Fig. 8B that this is not a major issue.

## Appendix C Stochastic simulations confirm that the steady-state assumption for ligand-receptor dynamics is justifiable at relevant cell speeds

Our analytical expression for a cell’s signal-to-noise ratio in Section II E assumes that the probability distribution of receptor binding is at steady-state given the local concentration of ligand. This assumption will be reasonable only if the cell moves to new concentrations sufficiently slowly, allowing the receptor binding statistics to equilibrate. How reasonable is this assumption? Ref. [17] found average speeds of single lymphocytes ranging from roughly 2 to 12 µm*/*min (Fig. 4F of [17]). Since the relaxation time for receptor equilibration for our chosen parameters is 10-15 minutes for receptor-internalizing cells (Fig. 1B) the fastest cells may move many cell diameters before receptor kinetics relax to equilibrium.

To determine the degree to which non-equilibrium receptor dynamics affect cell directionality, we performed Monte Carlo simulations of cells moving along a one dimensional track.

### 1. Stochastic receptor simulations with moving cells

We modeled the cell as a pair of compartments, each containing 25,000 receptors (see Fig. 2). As in Appendix A, all of the receptors are initially unbound. Unbound receptors can transform to bound receptors with a probability of *k*_on_[L]Δ*t* at each time step; bound receptors can transform to unbound receptors with a probability of *k*_off_Δ*t* or internalized receptors with a probability of *k*_in_Δ*t* at each time step; internalized receptors can transform to unbound receptors with a probability of *k*_rec_Δ*t* at each time step. Each receptor stays within its original compartment. The cell moves instantaneously with speed *v* in the direction of increasing *R*, i.e. at each timestep the position updates as *y*_i+1_ = *y*_i_ + sign(ΔR) *v*Δ*t*, where *v* is the cell’s speed and ΔR is the difference in R between the two compartments (cells for which ΔR = 0 choose a random direction at that time point). As in Appendix B, R is calculated at each time step and for each compartment from the fraction of bound receptors in each compartment.

In the experiments, cells attached to the track prior to CCL19 flowing into the chamber. Videos were taken three hours after CCL19 was initially flowed into the chamber. To mimic this setup, we held each simulated cell fixed at a randomly chosen position in the track’s concentration gradient (or outside of the track at constant concentration; see below) for three hours – i.e. we allow three hours for the cells to relax from the initial arbitrary choice of having all receptors unbound on the cell surface. We then allowed the cell to move for two hours with speed *v* in the direction of higher ΔR, as described above. We measured ΔR for the cell at each position it visited for the two hours during which the cell was moving. To measure the SNR in these simulated experiments, we divided the track into bins 26 µm long and placed each measurement of ΔR for all cells into its corresponding bin. To calculate the SNR at each bin, we took the average and sample standard deviation of all of the measurements ΔR recorded in that bin.

### 2. Boundary conditions and potential biases

Experimentalists and modelers trying similar analyses should note that we found that it is fairly easy for this sort of approach to develop surprising biases due to selection effects and boundary conditions. For instance, we initially tried assuming that cells could never be initialized outside of or re-enter the −500 µm+*ℓ* to 500 µm −*ℓ* region, where *ℓ* is half the length of the cell (Fig. 2B) However, this created clear biases. Even in the absence of a concentration gradient, with these boundary conditions, we found that the measured SNR for cells moving along a track with a constant concentration is negative at the left hand side and positive at the right hand side. This is from selection bias – cells that reach *y* = −500 µm + *ℓ* must have come from *y >* −500 µm + *ℓ*, and will by necessity have negative ΔR. A similar argument shows that cells at the other end of the track would have positive ΔR even in the absence of a concentration gradient. In contrast, cells can reach the middle of the track with either a positive or negative velocity. To alleviate these biases, we initialize cells from y = (500 µm + 120 min × *v* − *ℓ*) to y = 500 µm + 120 min × *v* − *ℓ*. 120 min × *v* is the maximum distance a cell can travel in the simulated experiment, and so this choice ensures that any point for which both compartments of the cell are within the real track is accessible by an equal number of cells on either side of that point. For this extension of the track, we set the ligand concentration for *y <* −500 µm to match the ligand concentration at *y* = −500 µm and to match *y* = 500 µm for *y >* 500 µm. For example, for the 0 to 25 ng/mL gradient case, [L] = 0 nM for all *y <* −500 µm and [L] = 2.3 nM = 25 ng/mL for all *y >* 500 µm. While our choice here is to minimize the effects of boundary conditions, it is likely that an attempt to experimentally measure SNR or chemotactic directionality as a function of *y* would also potentially be affected by these biases. We recommend future experiment-theory work take careful consideration of the true boundary conditions and check against experiments with no chemoattractant.

### 3. Qualitative agreement between stochastic simulation and simpler theory, with some caveats

Does our analysis of SNR in our stochastic model with a moving cell recapitulate the analytical results in the main paper, and maintain the simpler model’s agreement with the experiment? Fig 9 shows SNR_R_ from the binning analysis for our five primary cases of interest. We first check to see if our analysis quantitatively recaptures SNR as a function of *y* in the limit when the cell is motionless – *v* = 0 µm/min. In general, we see excellent agreement between the *v* = 0 stochastic model and the analytic theory, as expected. Note that for non-internalizing cells in the 0 to 500 ng/mL gradient the simulation results for the *v* = 0 µm/min case diverge slightly from the analytical result; this is because non-internalizing receptors have a significantly higher relaxation time, and so cells’ receptors in this case are further from equilibrium at the start of their trajectories – the 180 minute initial equilibration is not sufficient.

In general the measured SNR of cells moving at slower speeds like 3 µm/min agrees reasonably with the analytical result (Fig. 9). (This is the range relevant for [17], where speeds are most typically ~3 µm*/*min 7 µm*/*min, and rarely above 10 µm*/*min.) Simulated SNRs diverge from analytical theory notably at the edges of the track; we discuss this phenomenon more below. Even noting these differences, we see that the core qualitative results of the model in comparison to the experiment are preserved – the SNR is small in the 0 to 25 ng/mL gradient and the 400 to 500 ng/mL gradient, relevant and positive in the 0 to 100 ng/mL gradient, primarily chemorepulsive in the 0 to 500 ng/mL gradient for cells with receptor internalization, and chemoattractive in the 0 to 500 ng/mL gradient for cells without receptor internalization.

The measured SNR diverges from the analytical result at the edges of the track, and in some cases flips sign entirely. We believe that this effect again arises from a selection bias – because we track cells over time, the SNR_R_ of cells that accumulate at a given position *y* are not representative of the behavior of a cell initialized at that position. For example, if we look at the low concentration end of the 0 to 25 ng/mL gradient, the SNR is negative for non-zero velocities, even though the analytical result predicts a positive SNR for *y* ≈ −500 µm. For a cell to end up at *y* = −500 µm, it must either have traveled from *y <* −500 µm, where there is no chemoattractant, or from larger *y*. A cell that starts at *y* = −500 µm will quickly have a few ligands bound on its +*y* side and leave this end of the track. By contrast, a cell could also reach *y* = −500 µm by traveling from larger *y* if, by coincidence, the cell had more ligands bound on its lower-*y* side. Because the receptors take a time ~ *τ* to come to steady state, the ΔR for the cell will be negative when it reaches *y* = −500 µm. In other words, erroneously-moving cells are ‘attracted’ to regions that ‘repulse’ accurately-moving cells. The scale over which this effect reaches will depend on the cell’s speed *v* – we see much more dramatic effects at the highest speeds in the 0-25 ng/mL case in Fig. 9. We similarly see that at the “repulsive” point in the 0-500 ng/mL experiment in Fig. 9 that this repulsion is smoothed. As we would expect, the effects of finite speed are also much more dramatic in the 0-500 ng/mL experiment without internalization, when the correlation time *τ* is larger.

We see from our results in this Appendix that an attempt to experimentally measure SNR_R_ may be somewhat involved. Because of selection effects, binning the SNR_R_– or other quantifications of cell migration, including the velocity or directionality – will necessarily depend on both the chemoattractant gradient and on the assumptions made for how the cell interprets the signal to choose its direction. For instance, we have assumed that the cell always moves in the direction of higher R – including a tendency for the cell to move persistently would change these results.

